# Dew-induced transpiration suppression impacts the water and isotope balances of *Colocasia* leaves

**DOI:** 10.1101/178293

**Authors:** Cynthia Gerlein-Safdi, Paul P.G. Gauthier, Kelly K. Caylor

## Abstract

Foliar uptake of water from the surface of leaves is common when rainfall is scarce and non-meteoric water such as dew or fog is more abundant. However, many species in more mesic environments have hydrophobic leaves that do not allow the plant to uptake water. Unlike foliar uptake, all species can benefit from dew- or fog-induced transpiration suppression, but despite its ubiquity, transpiration suppression has so far never been quantified. Here, we investigate the effect of dew-induced transpiration suppression on the water balance and the isotope composition of leaves via a series of experiments. Characteristically hydrophobic leaves of a tropical plant, *Colocasia esculenta*, are misted with isotopically enriched water to reproduce dew deposition. This species does not uptake water from the surface of its leaves. We measure leaf water isotopes and water potential and find that misted leaves exhibit a higher water potential (*p* < 0.05) and a more depleted water isotope composition than dry leaves (*p* < 0.001), suggesting a ~30% decrease in transpiration rate (*p* < 0.001) compared to control leaves. We propose three possible mechanisms governing the interaction of water droplets with leaf energy balance: increase in albedo from the presence of dew droplets, decrease in leaf temperature from the evaporation of dew, and local decrease in vapor pressure deficit. Comparing previous studies on foliar uptake to our results, we conclude that transpiration suppression has an effect of similar amplitude, yet opposite sign to foliar uptake on leaf water isotopes.

## 1 Introduction

Non-meteoric water (NMW, dew or fog) is an important source of water for many plants that occurs consistently in all environments (Burgess & Dawson, 2004; Breshears et al., 2008; Lakatos et al., 2012; Berry, Hughes, & Smith, 2014; Hill et al., 2015). Specifically, dew is the condensation of atmospheric water vapor on a surface, usually at night when the surface is cooling down to below the dew point temperature of the air (Beysens, 1995). In the case of fog, the condensation happens around particles in suspension in the air, such as soot, sea salt, or organic matter (Price & Clark, 2014). For this reason, fog water can be transported and its isotope and chemical composition can be very different to the atmospheric water vapor at the place of collection, whereas dew water is usually similar to that of local atmospheric water vapor composition. The use of NMW has been reported for all types organisms, from trees to crops (Wang et al., 2016), and the evolutionary adaptations that plants have developed to utilize NMW are diverse and creative (Aparecido et al., 2017), from water trapping trichomes (Kim et al., 2017) to enlisting the help of fungal hyphae (Burgess & Dawson, 2004).

The water supplied by NMW can be directly uptaken by the leaves, a phenomenon called foliar uptake (Yates & Hutley, 1995; Eller et al., 2013; Berry, White, & Smith, 2014; Oliveira et al., 2014; Baguskas et al., 2017). Evidence for foliar uptake of fog water has been mainly provided by the comparison of the stable isotope composition of the leaf water and of the NMW (Limm et al., 2009; Gotsch et al., 2013; Berry, Hughes, & Smith, 2014; Goldsmith et al., 2017). Indeed, different pools of water in the ecosystem will have very distinct isotope compositions (Kaseke et al., 2017) which can be distinguished even after the water enters the leaf through foliar uptake. For example, water that undergoes evaporation, like water contained in shallow soils, will be more enriched in heavy isotopes than rain- or groundwater. For the same reason, water inside leaves is usually more enriched than the water contained by stems or roots.

NMW can also interact with the leaf energy balance, without actually entering the organism. For example, dew can reduce incoming radiation through the increase in albedo due to the presence of water droplets at the surface of the leaf. In addition, the energy subsidies created by the evaporation of the water droplets will decrease leaf temperature. Both processes will lead to a decrease in leaf transpiration (Tolk et al., 1995). Transpiration suppression is often mentioned as an existing response mechanism of vegetation to NMW deposition (Berkelhammer et al., 2013; McLaughlin et al., 2017) but its effect have so far not been quantified. On the one hand, transpiration suppression from NMW deposition will impact the expected leaf water isotope composition. In particular, decreasing leaf transpiration caused by a decrease in incoming energy to the leaf should lead to leaf water depletion in heavy isotopes (Farquhar & Cernusak, 2005; Cernusak & Kahmen, 2013). On the other hand, NMW is usually enriched in heavy isotopes compared to rain water (Scholl et al., 2010), which leads to a clearly enriched signal in leaves that use foliar uptake. The impact of transpiration suppression on the isotope composition is therefore likely to be opposite to that of foliar uptake, but foliar uptake studies have so far not taken transpiration suppression into account, even though it likely results in an underestimation of the amount of water taken up by the leaf or transpired.

The objective of our study is to quantify the impacts of NMW on the coupled leaf water and energy balance, a phenomenon that has so far only been mentioned in previous work (Dawson, 1998; Limm et al., 2009; Berkelhammer et al., 2013). Here, the effects of dew deposition on the leaf water potential, transpiration rate, and water stable isotopes are experimentally determined. To dissociate any observed effect from foliar uptake, isotopically-enriched dew is used, and the experiments are conducted on *Colocasia esculenta*. This species is native to South East Asian tropical forests but has been cultivated across the world for many centuries under the name of taro. With a contact angle of ~164°(Neinhuis & Barthlott, 1997), *Colocasia esculenta* is considered to have highly water-repellent leaves, which can reach a size of up to c. 50 cm in length and c. 40 cm in width. It is a very unique plant, with amphistomatic leaves that are adapted to shaded environments (Onwueme & Johnston, 2000) and occasional flooding (Mabhaudhi et al., 2013). This species was specifically chosen because of it does not have the capacity to uptake water through foliar uptake.

We apply a protocol using the Picarro Induction Module (IM) coupled to a cavity ringdown spectrometer for the fast analysis of small-sized leaf samples, allowing for spatial and temporal high-resolution mapping of leaf water isotopes (Gerlein-Safdi et al., 2017). We then analyze the spatial patterns of leaf water isotopic enrichment of leaves that have been sprayed with isotopically enriched water, simulating dew deposition. We show that dew deposition decreases transpiration and increases water potential in *Colocasia esculenta* leaves, and we discuss three possible mechanisms to explain this result. Finally, we compare our findings to previous studies focused on foliar uptake, leading us to the conclusion that transpiration suppression and foliar uptake have an opposite and comparable effect on leaf water isotopes.

## 2 Materials and Methods

### 2.1 Laboratory experiment

Our first experiment examines leaf scale spatial and temporal patterns of water isotopes induced by the presence or the absence of dew. The sampling and analysis closely follow the method described in Gerlein-Safdi et al.(2017). A plant of *C. esculenta* was planted in a 57-liter (~ 15 gallons) pot filled with garden soil (Miracle Gro, Marysville, OH, USA) and grown to maturity. The natural patterns of *Colocasia esculenta* water isotope have been presented in details in a previous study (Gerlein-Safdi et al., 2017). The plant was watered daily with tap water (*δ*^18^O ≃ −6.0 ‰, *δ*^2^H ≃ −38 ‰) for multiple weeks.

Two leaves of c. 30 cm length and of the same *Colocasia esculenta* plant were cut at the junction of the petiole and the rachis and placed c. 80 cm under a light (Eiko 1960 EBW, 500 W, 10500 lumens, color temperature of 4800 K). The entire experiment lasted four hours. During that time, the adaxial side of the treated leaf was misted with isotopically-labelled water (*δ*^18^O ≃ 8.8 ‰, *δ*D ≃ 737 ‰) every half-hour, while the control leaf was left untouched during the entire experiment. After four hours, any residual water was dried by gently padding the leaf with a paper towel and samples were collected from both leaves as described in Section 2.2.

In our second experiment, we focused on the effect of water droplet deposition on leaf water potential under high water stressed conditions. Leaves were cut at the junction of the petiole and the rachis and left to desiccate on the lab bench. Three different water stress conditions were tested: natural drying (control), high heat drying, and high heat and mist. In the high heat case, the leaf was placed 80 cm under a light as specified above and the desiccation tracked for about eight hours. In the high heat and mist case, the leaf was also misted with ultra pure water every hour using a spray bottle. Again, surplus water was allowed to runoff, leaving the leaf covered in submillimeter size water droplets. Leaf disks of 2.5 cm diameter were collected every hour, and immediately weighted. As recommended by the instrument manufacturer, the surface of each leaf disk was wetted with ultra pure water, sanded with ultra-fine sandpaper (3M, 600 grit sandpaper), and the water potential analyzed on a dew point potentiometer (WP4C by Decagon Devices Inc.).

### 2.2 Isotope analysis

For the water isotope analysis, leaf samples were analyzed using an Induction Module (IM) combined to a Cavity Ring Down Spectrometer (CRDS) L2103-i from Picarro Inc. (Sunnyvale, CA, USA). Each leaf was sampled in 16 different locations. All of the sampling points were located on the same half of the leaf and each point consisted of four holes (6 mm diameter) punched next to each other forming a square. Each hole was punched as quickly as possible to avoid evaporation, which would influence the isotope composition of the neighboring holes. Leaf disks were individually secured in aluminum strips provided by the IM manufacturer and inserted in a 4 mL sealed glass vial provided by the instrument manufacturer. It took about 15 min to sample a single half-leaf. The analysis started as soon as the sampling was over. The prepared vials were then stored in the fridge until being analyzed.

#### 2.2.1 IM-CRDS analysis sequence

The IM was set on the ‘normal leaf’ setting, which has been shown to dry the samples completely without burning them (Gerlein-Safdi et al., 2017). The IM was equipped with a micro-combustion module (MCM) to reduce the interferences due to the presence of organics (Dennis et al., 2014) in water samples extracted from plants (A. G. West et al., 2010; Chang et al., 2016). Each half-leaf was sampled in 16 different locations, which corresponds to 64 punched holes per half-leaf and a sampling density of c. 6.5 samples per dm^2^. The IM analysis lasted c. 1.5 day per half-leaf.

The IM-CRDS analysis sequence was adapted from a protocol developed in van Geldern & Barth (2012) for liquid water samples and is described in details in Gerlein-Safdi et al. (2017). Six empty vials were run at the beginning of each analysis. The average water vapor content, *δ*^18^O, and *δ*^2^H of the six vials were measured and introduced in a mixing model that allowed the signal from the ambient air present in the vial to be removed to retrieve the true isotope composition of the sample analyzed:

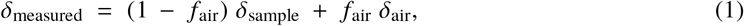

where *f*_air_ = [H_2_O]_air_ / [H_2_O]_measured_, with [H_2_O]_air_ and [H_2_O]_measured_ the water vapor concentrations inside the blank vial and the sample vial, respectively. Reference water samples (ABB - Los Gatos Research Inc., San Jose, CA, working standards, all vs VSMOW: ER4: *δ*^18^O = 81.33±0.3‰, *δ*^2^H = 654.1±1.0‰, ER1: *δ*^18^O = 12.34±0.3V *δ*^2^H = 108.7±1.0‰, 5A: *δ*^18^O = -2.80±0.15‰, *δ*^2^H = -9.5±0.5‰, and 3: *δ*^18^O = −11.54±0.1‰, *δ*^2^H = -79.0±1.0‰) were injected on punch holes of glass filter paper provided with the IM and the same piece of filter paper was reused for all the injections of a single reference water. We found that 3*μ*L of reference water were necessary to reproduce the peak of water vapor concentration (in the analyzer) produced by one punch hole of *C. esculenta*(Cui et al., 2017).

Following the protocol developed for liquid water samples in previous work (van Geldern & Barth, 2012), the data were corrected for drift and memory effects, and rescaled back to VSMOW. The drift of the instrument was determined by rerunning the drift-monitoring standard (DEST, working standard ER1) at the beginning, the end, and half-way through the samples. This standard water was chosen to have an isotope composition as close as possible to that of the leaf samples analyzed (average leaf isotope composition: *δ*^18^O = 55.1‰, *δ*^2^H = 132‰). A linear regression was then used to realign all the measurements, according to the sample order. The effect of the drift was found to be small for both isotope ratios, with a slope of −0.004±0.005 ‰ per sample in *δ*^18^O and −0.05±0.07 ‰ per sample in *δ*^2^H. These values are comparable to values found for liquid water samples (van Geldern & Barth, 2012). For the memory effect, three standard waters with large isotopic differences were injected 10 times in a row each. An isotopically high (HIS, working standard ER4) and low (ANTA, working standard 3) standard waters were used along with DEST to determine memory effects from high-to-low and low-to-high transitions. A memory coefficient was associated with each successive injection. The coefficients were determined using the ‘Solver’ function in Excel (Microsoft, Redmond, WA, USA) so as to minimize the standard deviation of all three standard waters simultaneously. As shown in Gerlein-Safdi et al. (2017), the evolution of the memory coefficients has a similar exponential shape for all our runs, and the memory effect has completely disappeared by the fifth injection. The scaling to VSMOW was done using the DEST, HIS, and ANTA standard waters of known isotope composition and using a linear regression to correct for potential deviations from the 1:1 line. A fourth intermediate standard water (HERA, working standard 5A) was used for quality control and we found a precision of ±0.2‰ (SE of the difference from true value) in *δ*^18^O and ±1.1‰ in *δ*^2^H.

#### 2.2.2 Isotope mapping

Once the sampling was completed, the leaves were pressed and scanned. We digitally located the positions of the sampling locations, the midrib, and the secondary veins. To interpolate the isotope composition of the leaf in-between sampling locations, we used an inverse distance weighting (IDW) method with a fixed weight parameter available in the ‘gIDW’ add-on package developed by Giuliano Langella for MATLAB (Mathworks Inc., Natick, MA, USA). The IDW method was specifically chosen because it is commonly employed for isotope mapping (J. B. West et al., 2009).

### 2.3 Environmental and physiological data

Stomatal conductance, *g*_s_, relative humidity, *h*, and leaf and air temperatures, *T*_leaf_ and *T*_air_, were recorded throughout the growth of the plant. Air temperature and relative humidity in the lab were remarkably stable throughout the growth with *T*_air_ = 26.3±1.2 °C and h = 41.2±4.4 % (average ± STD). Stomatal conductance and leaf temperature were recorded during growth using an SC-1 Leaf Porometer (Decagon Devices Inc., Pullman, WA, USA). Measurements were taken every other day at 11 locations around the leaf and on the abaxial and the adaxial sides of the leaves. Six measurements were taken at each locations and the average taken to obtain a value at one location, on one side and on one day. Average stomatal conductance was 0.0485±0.08 mol m^-2^ s^-1^ and leaf temperature was 26.2±1.5 °C. Stomatal conductance was not available during the experiments. Leaf temperature during the experiments was monitored every half-hour using an infrared camera (Flir T500-Series, Wilsonville, OR, USA).

### 2.4 Linking d-excess and transpiration

While d-excess is commonly used in Atmospheric Science (Risi et al., 2013) and for interpreting ice core data (Luz et al., 2009), it has not been widely used in plant physiology. However, because it combines both ^2^H and ^18^O, d-excess contains more information than the isotopologues taken separately. Indeed, lower (more negative) d-excess values are associated with higher transpiration rates (Voelker et al., 2014). To interpret d-excess differences in terms of transpiration rates, we link d-excess to steady-state relative humidity. The steady-state enrichment of leaf water Δ*_E_* above source water is expressed in (Farquhar et al., 2006) as

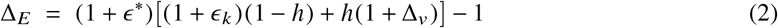

where *h* is the relative humidity, *ϵ*_*_ is the equilibrium fractionation; *ϵ*_*_ = 9.2 *‰* (74 ‰) for ^1^H_2_^18^O (^1^H^2^HO) at 25°C (Craig & Gordon, 1965). The kinetic fractionation factor, *ϵ_k_*, is taken as

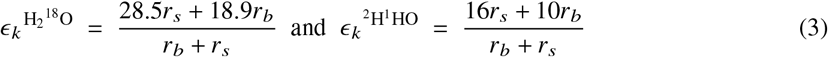

for ^1^H_2_^18^O and ^2^H^1^HO, respectively (Farquhar et al., 1989, 2006). *r_s_* is the stomatal resistance and it is taken to be constant and equal to 840 s m^-2^ (based on the value described in Section 2.3). The resistance of the boundary layer, *r_b_*, depends on leaf size and wind speed. Here we choose a constant leaf size of 40 cm and a wind speed of 0.2 m s^-1^, resulting in an *r_b_* = 1.13 10^5^ s m^-2^. Δ*_v_* is the enrichment of ambient water vapor above source water, which was calculated for a measured air composition of *δ*^18^O = −17 ‰ and *δ*^2^H = −100 ‰ and a source water corresponding to the tap water used to water the plants (*δ*^18^O ≃ −6.0 ‰, *δ*^2^H ≃ −38 ‰).

Δ*_i_*, the enrichment of a sample *i* relative to a source can be linked back to isotope compositions expressed in *δ* notation through the relative ratios *R*:

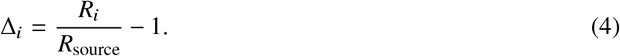

Finally, we can express *R_i_* as a function of *δ_i_*, and obtain a relation between *Δ_i_* and *δ_i_*:

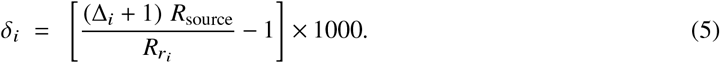

By replacing Δ*_i_* in Equation 5 by its expression from Equation 2 and combining the expressions for *δ*^18^O and *δ*^2^H, we obtain an expression for the d-excess as a function of the relative humidity h. We solve for h, bounding its value between 0 and 1. Assuming that the vapor pressure inside the leaves, *e_i_*, is at saturation, we may then calculate the estimated transpiration rate E (in mmol m^-2^ s^-1^) as

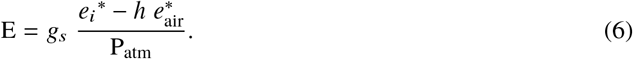

Here, P_atm_ is the atmospheric pressure taken to be 101.3 kPa and *g_s_* (in mmol m^-2^ s^-1^) is the stomatal conductance equal to 1/*r_s_*. *e_i_*^*^ (in kPa) is the saturated vapor pressure calculated for *T*_leaf_ of 25°C. 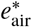 (in kPa) is the saturated vapor pressure at *T*_air_, which is also taken to be 25°C. In our analysis, we compare the transpiration rates of misted leaves, E_dew_, and that of control leaves, E_control_.

### 2.5 Competing effects of foliar uptake and transpiration suppression

To compare the relative effects of foliar uptake and transpiration suppression on leaf water isotopes, we analyzed the results of three different studies that conducted similar experiments on different species. Even though these studies looked at foliar uptake of fog, we chose them because they conducted identical experiments and covered a broad range of species. Limm et al. (2009) looked at a ten different species from the coastal redwood ecosystem of California (*Pseudotsuga menziesii* and *Sequoia sempervirens* (conifers), *Polystichum munitum* and *Polystichum californicum* (ferns), *Oxalis oregana* (a short herbaceous), *Arbutus menziesii, Gaultheria shallon, Vaccinium ovatum, Notholithocarpus densiflorus* and *Umbellularia californica* (all evergreen broadleaf)), while Eller et al. (2013) focused on *Drimys brasiliensis*, a woody broadleaf evergreen native from Central and South America, and Berry & Smith (2014) concentrated on *Abies fraseri* and *Picea rubens*, two montane conifers from the Appalachian Mountains. All the studies conducted greenhouse experiments in which saplings experienced nighttime fog. Leaf samples were collected in the evening before the fogging treatment and in the morning, right after the treatment. Every study used isotopically labeled fog with a different composition (*δ*^2^H_fog_ − *δ*^2^H_soil_ = 16 ‰ in Berry & Smith (2014), 78 ‰ in Limm et al. (2009) and 712 ‰ in Eller et al. (2013)). To compare the different experiments, we normalized the results of the three experiments to reflect the effect of a fog water enrichment of 20 ‰ above soil water composition, since this is within the range of natural values (Scholl et al., 2010).

### 2.6 Statistical analysis

Responses for the different experiments were analyzed using a two-sample t-test (Welch’s t-test) with a 5% significance level. This test has been recognized as a better alternative to the Student’s t-test when dealing with groups of unequal sample size or variance (Ruxton, 2006). In the following, we will report the p-value, p, the test statistics, t, and the degrees of freedom of the test, v. When comparing the results of the different treatments, we treated the multiple samples collected on each leaf as a single population. In the following, *stat* and *syst* refer to the statistical and the systematic errors, respectively.

## 3 Results

### 3.1 Water isotopes

The results of Experiment 1 are presented as maps of the analyzed half leaves (Figure 1). In this case, the lamp artificially increases the transpiration rate in both the control and the misted leaves, leading to significantly enriched *δ*^18^O and *δ*^2^H values and low d-excess values. The d-excess in the control case is c. 173.0 ‰ more negative than for the misted leaves (two-sample t-test: t = 3.9, *ν* = 29, *p* < 0.001).

The misted leaf in Experiment 1 is less enriched in heavy isotopes than the control, despite being misted with highly enriched water (*δ*^18^O ≃ 8.8 ‰, *δ*^2^H ≃ 738 ‰). In addition, the control leaf shows significantly lower d-excess values than the misted leaf. As explained in Section 2.4, d-excess is a measure of how evaporated a pool of water is. More negative values in the control leaf are consistent with more transpiration than in the misted leaf. This is an indication that the misting decreased transpiration in the misted leaf.

**Figure 1:**
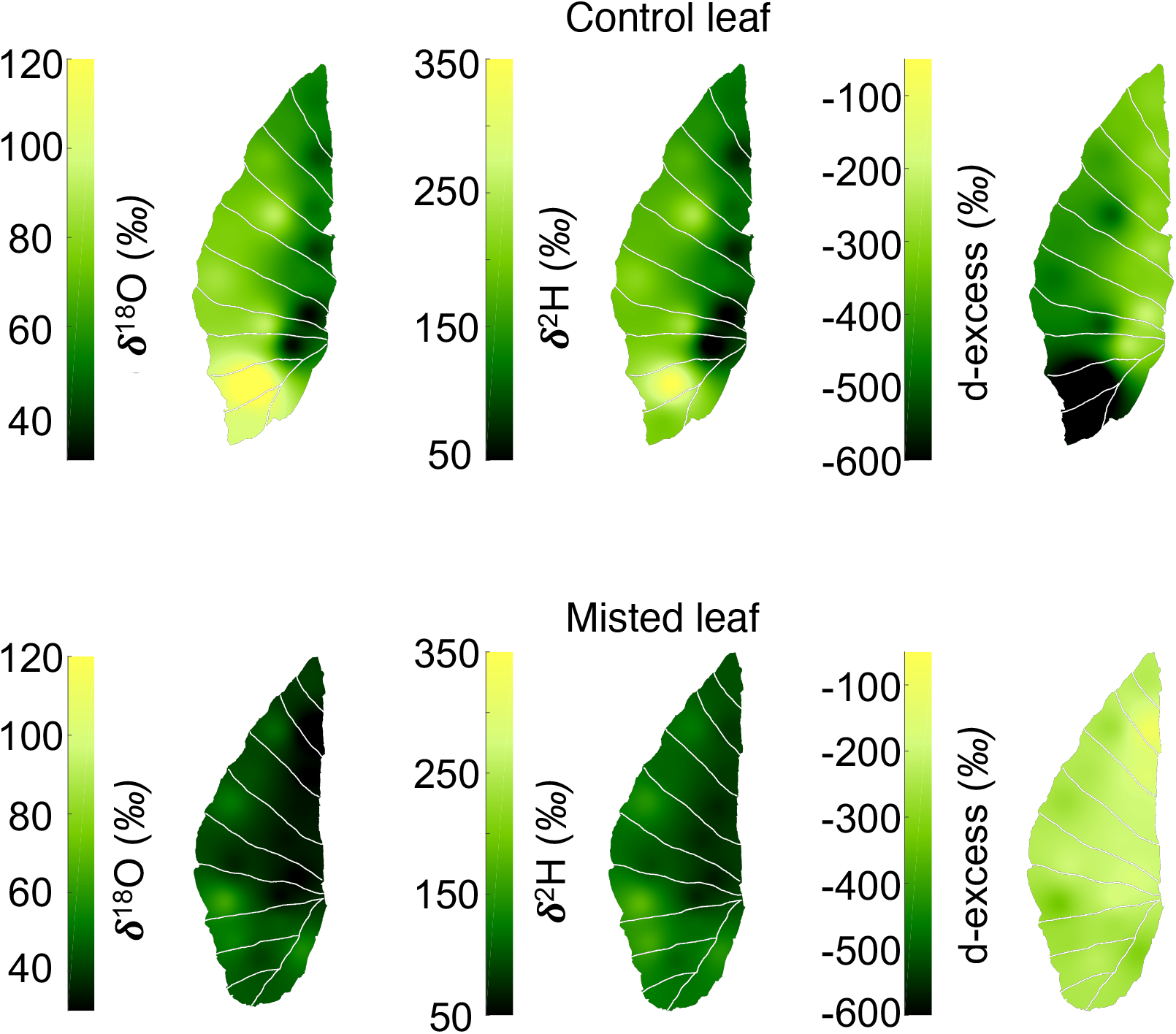
Maps of two leaves left under a 500W light for four hours. **Top row:** δ^18^ O, δ^2^ H and d-excess of the control (not misted) leaf. **Bottom row:** δ^18^ O, δ^2^H and d-excess of the leaf misted with isotopically enriched water(δ^18^ O ≃ 8.8 ‰, δ^2^ H ≃ 738 ‰) every half-hour. The control leaf shows higher enrichment and lower d-excess values that are associated with enhanced transpiration compared to the misted leaf.

We apply the model described in Section 2.4 to interpret our results in terms of differences in transpiration rate and find that the mist treatment significantly (t = − 3.9, *μ* = 29, *p* < 0.0001) decreases transpiration by 29.9 ±9.1 (stat) %. These values are consistent with (Garratt & Segal, 1988), who estimated that the reduction in transpiration due to dewfall could reduce daily plant water use by c. 8%. This value was obtained for wheat plants associated with a low transpiration rate. Because *Colocasia esculenta* leaves are larger and have a higher transpiration rate, we expect the reduction to be larger in our case (see Section 4.1).

The use of a constant stomatal conductance and leaf temperature are most likely the largest sources of systematic error in this estimate of transpiration differences. To calculate the systematic error associated with our choices for those parameters, we calculated E_dew_/E_control_ for a range of leaf temperatures from 10° to 40°C and for a stomatal conductance *g_s_* from 0.1 to 0.8 mmol m^-2^ s^-1^. We found that the systematic error associated with our choice of *g_s_* and *T*_leaf_ were both negligible.

### 3.2 Leaf water potential

Experiment 2 looks at the temporal evolution of water potential in desiccating leaves (Figure 2). The pressure-volume curve (Figure 2a) shows a shape characteristic of a high apoplastic fraction (Bartlett et al.,2012). Indeed, *Colocasia esculenta* is known for its high content of mucilage (Quach et al., 2001; Njintang et al., 2014). The polysaccharides constituting the mucilage have been shown to have a large impact on leaf water potential and plant resistance to water-stress (Morse, 1990): because of their high capacitance (they can hold more than ten times their weight of water), the polysaccharides create a large apoplastic capacitor of available water that can buffer the changes in water potential associated with large variations in leaf water content.

**Figure 2:**
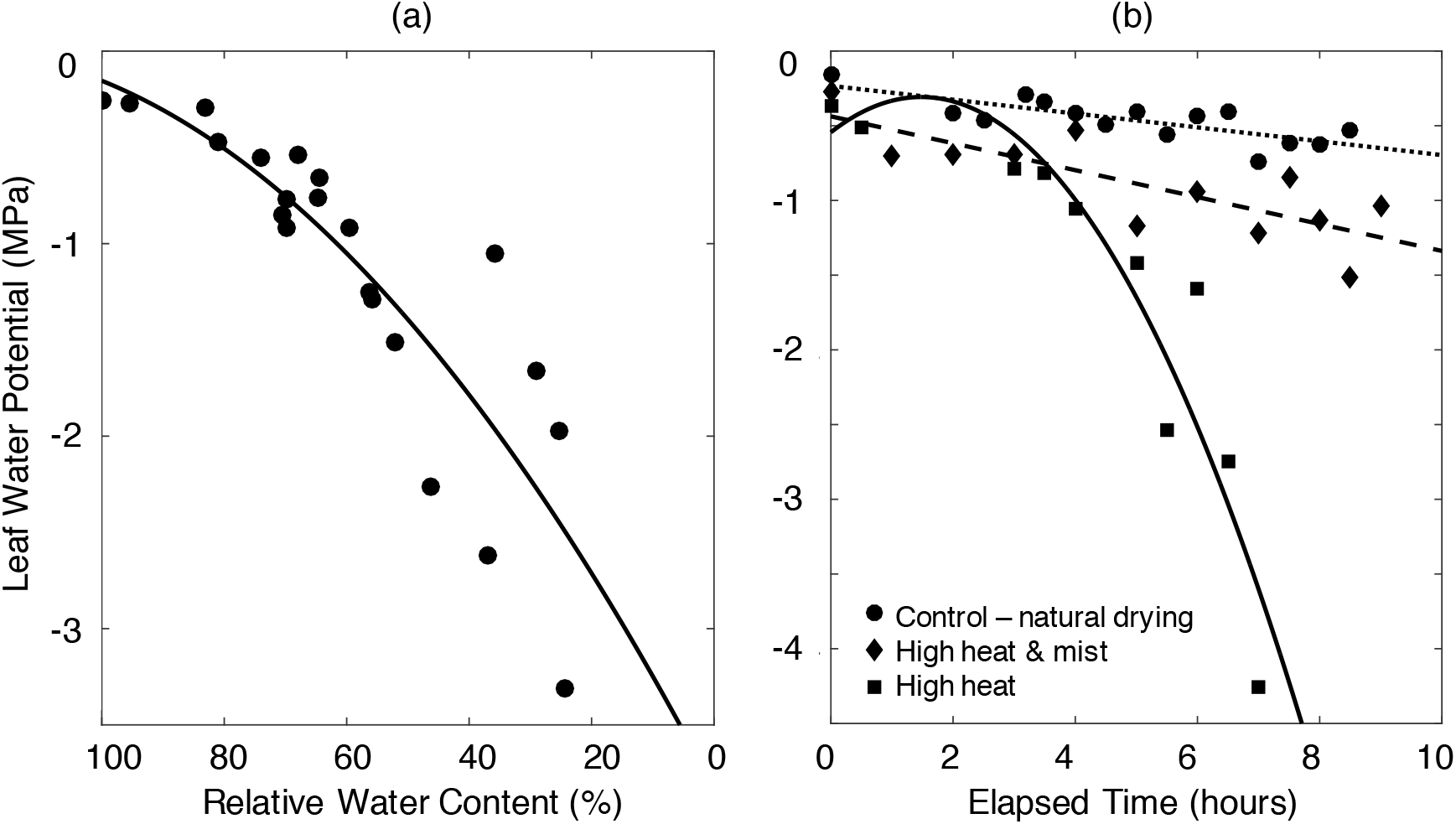
Desiccation curves of Colocasia esculenta leaves. **(a)** Pressure-volume curve for Colocasia esculenta. The shape of the curve is typical characteristic of a high apoplastic fraction (Morse, 1990; Bartlett et al., 2012). The black line represent a second-order polynomial fit. This plot combines data from two leaves of the ‘high heat and mist’ treatment, in order to represent the whole range of leaf water potentials observed during the experiment. **(b)** Typical examples of the temporal evolution of the leaf water potential of Colocasia esculenta leaves under three different treatments. All the leaves under the natural drying (circles) and the high heat and mist (diamonds) treatments are well fit by a linear relation (dotted and dashed lines, respectively). All but one of the leaves under the high heat drying case (squares) are better fit by a parabola (solid line). All the leaves shown here are c. 38 cm long.

For the control and the high heat and mist cases, the leaf water potential experiences a slow decline, which is well approximated by a linear function (Figure 2b). However, the high heat treated leaves experience a faster decline and are better approximated by a parabola. Table 1 presents the average decline from initial to final leaf water potential for the three different treatments. All the data is normalized for leaf size and drying time. The decline in water potential was c. 64% smaller in the misted leaves than in the leaves subjected to the same high heat treatment but that did not get misted (two-sample t-test: t = 2.37, *ν* = 7, *p* < 0.05). The decline observed for misted leaves is not statistically different to the one observed for naturally drying leaves (two-sample t-test: t = −1.46, *ν* = 6, *p* = 0.19).

**Table 1.** Average decrease in water potential (MPa) for the three treatments of Experiment 2: ‘Natural drying’ (control), ‘High heat and mist’ and ‘High heat’. Third column shows one standard error and n is the number of replicates for each treatment. All the data was normalized to reflect the drop in water potential for a 40 cm long leaf over 8 hours.

| Treatment | Average drop in leaf water potential over 8h (MPa) | SE | n |
| --- | --- | --- | --- |
| Control | 0.43 | 0.03 | 3 |
| High heat & mist | 1.05 | 0.31 | 5 |
| High heat | 2.9 | 0.77 | 4 |

### 3.3 Comparison to foliar uptake

Using previous studies on foliar uptake (described in Section 2.5), we were able to compare the relative impact of both processes. Foliar uptake has the largest impact on conifers (Figure 3), where the difference in enrichment between treatment and control reaches up to c. 20 ‰. Transpiration suppression from water deposition exhibits the opposite effect, with a magnitude similar or larger to the largest foliar uptake case. The three foliar experiments presented here all used nighttime treatment, so transpiration suppression did not impact the enrichment observed. However, the competing effects of foliar uptake and transpiration suppression are likely to be very important when analyzing field or day time foliar uptake experiment data.

**Figure 3:**
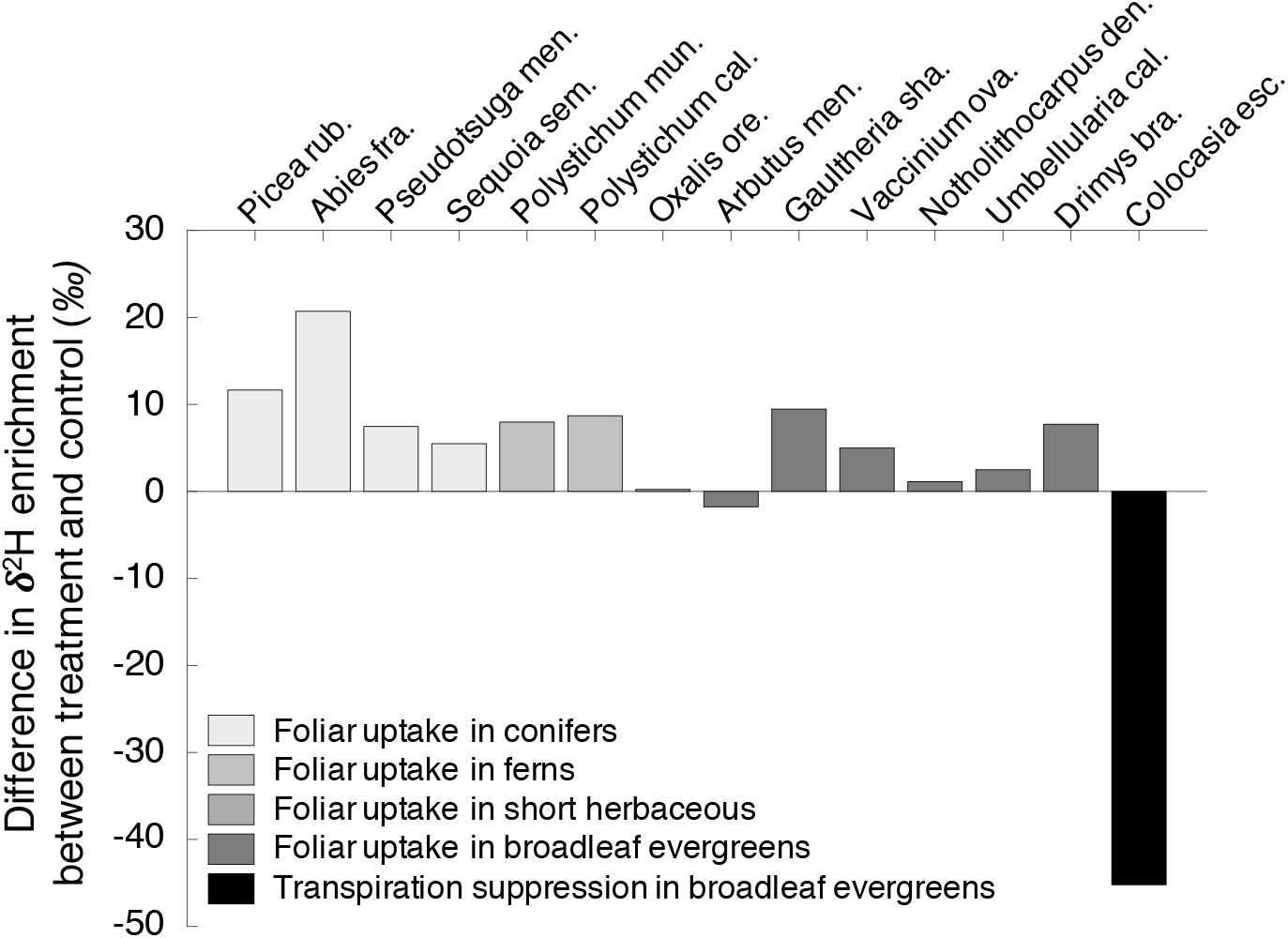
A comparison between the impact of foliar uptake of nighttime fog in three studies (Limm et al., 2009; Eller etal., 2013; Berry, White, & Smith, 2014) to transpiration suppression in Colocasia esculenta. Bars represent the magnitude of the difference in δ^2^H enrichment between misted and control plants. Here, enrichment is the difference between pre- and post-treatment leaves. All the foliar uptake data was normalized to reflect the enrichment corresponding to a realistic difference of 20 ‰ between the source water (watering water) and the fog water (Scholl et al., 2010).

## 4 Discussion

### 4.1 Leaf energy cycle

Our results show that the deposition of submillimeter size droplets on its surface allows the leaf to decrease its transpiration rate and maintain its water potential. The results presented in this study are consistent with a lack of foliar uptake on the adaxial side of *Colocasia esculenta* leaves. Indeed, if foliar uptake was taking place, we would expect to find that the misted leaf had leaf water enriched in heavy isotopes due to the presence of the enriched misted water inside the leaf. This is not the case and transpiration suppression is then the only phenomenon inducing differences in leaf water isotope composition between the treated and control leaves.

We found that the water balance of the leaf is influenced by the change in energy balance associated with the water droplets deposited at the surface and we identified three distinct processes that could lead to the observed effect.

First, the deposited droplets increase the albedo of the leaf, allowing more of the radiation to be reflected away from the leaf. Depending on the direction of the incoming solar radiation, water can have an albedo as high as 1 (perfect reflector) whereas typical values for leaves are c. 0.2. The increase of vegetation albedo due to dew deposition has been observed many times in the field (Pinter, 1986; Zhang et al., 2012). By reflecting more radiation when they are wet, leaves will decrease the incoming shortwave radiation and consequently keep their temperatures lower. In our experiment, we found a difference of c. 1.3°C between the ‘high heat and misted’ and the ‘high heat’ treatments (data not shown), confirming this hypotheses. This will in turn reduce the evaporative demand and the leaf transpiration.

Second, the energy that is not reflected will be dissipated through the evaporation of the droplets. Because evaporation is an exothermic process, the evaporation of the water droplets will result in the cooling of the leaf surface (Monteith, 1965). This will again reduce the evaporative demand and the transpiration.

Finally, the evaporation of the droplets will cause the air close to the leaf to have a higher relative humidity than the surrounding air (Defraeye et al., 2013), creating a moist micro-climate around the leaf (Jones, 1992). This will decrease the difference between the interstitial and the air vapor pressures, and reduce the flux of water vapor out of the leaf, namely transpiration. By decreasing the outward flow of water vapor, more CO_2_ will be able to enter the leaf, increasing interstitial CO_2_ concentration, photosynthesis, and water use efficiency. The increase in surface roughness associated with the presence of the droplets at the surface of the leaf will also contribute to increasing the size of the boundary layer. Water potential values are correlated with leaf relative water content (Maxwell & Redmann, 1978) and with stomatal conductance (Lhomme et al., 1998); by maintaining a higher water potential, the leaf will be able to open its stomata wider. CO_2_ assimilation is in turn linearly correlated to stomatal conductance (Lambers et al., 2008). As a result, by affecting the leaf energy cycle, dew deposition will allow the leaf to maintain its water status and increase CO2 assimilation through multiple mechanisms.

The three processes described above are not mutually exclusive and are happening simultaneously. We speculate that the cooling provided by the evaporation of the dew will have the largest impact on transpiration suppression. However, the balance between all three mechanisms will likely be species dependent. For example, in the specific case of *Colocasia esculenta*, because the leaves are amphistomatous we expect that the effects on the leaf boundary layer and CO_2_ uptake will likely be less important than for a hypostomatous leaf for which the dew is not clogging half of the stomata.

### 4.2 Implications for foliar uptake studies

By decreasing transpiration, dew deposition suppresses the isotopic enrichment associated with leaf water transpiration (Farquhar et al., 2006). Therefore, dew-wetted leaves will have a bulk isotope composition lower (more depleted in heavy isotopes) than leaves that do not experience it. The average *δ*^2^H enrichment difference between the misted and the control leaves reaches -45±35 ‰. The effect of dew is “artificially” increased by the high transpiration rate caused by the lamp, but it still gives a first order value for transpiration suppression. NMW is usually more enriched in deuterium than rain and soil water by up to 50 ‰ (Scholl etal., 2010; Kaseke et al., 2017). If foliar uptake is indeed happening in a leaf, the uptake of heavy fog or dew water will then enrich the leaf water, while transpiration suppression depletes leaf water in heavy isotopes. Limm et al. (2009)already pointed out how foliar uptake and reduced nighttime stomatal conductance due to the saturated atmosphere during fog events could have opposite effects on the leaf isotope composition. Daytime transpiration is a much larger water loss for plants than nighttime water vapor fluxes and the effects of dew deposition during day time is expected to be have an even larger impact on leaf isotopes than that discussed by Limm et al. (2009).

In addition, Berry, White, & Smith (2014) observed a significantly larger enrichment when fogging saplings in the morning than in the afternoon. This results is well explained if transpiration suppression is taken into account, because it will have a larger effect in the afternoon, when leaves are hotter and radiation stronger. Our results suggest that, in the field, transpiration suppression might have a larger impact on leaf isotopes than foliar uptake, although the relative importance of both effects will depend on many factors, including the isotope composition of the NMW (dew or fog), the timing and length of the wetting event, the size of the leaf, the atmospheric conditions, and of course, the species.

Finally, it is crucial to remember that while transpiration suppression and foliar uptake have opposite effects on the leaf water isotopes, both processes will increase leaf water content. Since both mechanisms are still poorly understood and highly species-dependent, it is difficult to provide an accurate estimate of the cumulated effect of foliar uptake and transpiration suppression. Limm et al. (2009) found a maximum increase in leaf water content of c. 12%. In our study, we found a c. 30% decrease in transpiration rate. If we take the example of a 0.05 m^2^ *Chrysanthemum morifolium* leaf with an absolute leaf water content of 20 mg (Hughes et al., 1970) and an average stomatal conductance to CO_2_ of 50 mmol m^-2^ s^-1^(Ozturk et al., 2013), which corresponds to a conductance of 80 mmol m^-2^ s^-1^ for water vapor. A first order estimate gives a daily flux of water vapor is c. 2300 mol m^-2^ for an 8 *h* day, equivalent to about 41.5 mm (or kg m^-2^) of water transpired per day. A 30% decrease in transpiration would amount to a total of 12.5 mm of water saved over a day, while fog might provide about 2.4 10^−6^ mm of water. It might seem that transpiration suppression has an overwhelmingly larger effect on leaf water than foliar uptake does, but the value for foliar uptake is likely underestimated, since it does not take into account that most water in the leaf is actually replaced by soil water throughout the day and that a 12% fog water inside the leaf likely corresponds to a lot more than simply 12% of the total leaf water content.

The results of our study show a large impact of transpiration suppression from water droplet deposition. The simultaneous occurrence of NMW deposition and drought is common in drylands (Agam & Berliner,2006; Kaseke et al., 2017), where many plants rely on NMW as their primary source of water (Stanton & Horn, 2013). Regular dew formation has also been observed in the upper canopy of the Amazon forest during the dry season (Satake & Hanado, 2004; Frolking et al., 2011) and leaf wetness can last almost all day in the understory of tropical forests (Aparecido et al., 2017). In these cases, the energy balance is thought to be the main driver of leaf water isotope composition, with a response much larger than to soil water availability, for example (Wayland, 2015). Transpiration suppression will delay the time when leaves reach their maximum transpiration rate and attain isotopic steady state (Dubbert et al., 2014). Isotopic steady state is often assumed when interpreting transpiration data, but Dubbert et al. (2013) recently showed that this assumption is typically unjustified and can lead to errors in estimated transpiration fluxes by up to 70%, because steady state models systematically overestimate the isotopic enrichment of leaf water. Isotopic steady state depends on the leaf transpiration rate, which changes quickly as energy flux incident on the leaf changes, for example when the leaf goes from the shade to the sun (Smith & Berry, 2013). The impact of NMW deposition has a large impact at short time scales on both leaf transpiration and water isotopes because of this fast response.

### 4.3 Potential limitations

One of the limitation of this study is the use of excised leaves. In Table 2, we show the average enrichment in heavy isotopes in the longitudinal and the radial direction. Previous work found that intact *Colocasia esculenta* leaves have a pronounced radial enrichment and no enrichment in the longitudinal direction, likely due to the major to minor vein ratio (Gerlein-Safdi et al., 2017). Here, we find that the hydraulics of the cut leaves still follow similar patterns of enrichment, with a significant increase in leaf water enrichment in the radial direction moving perpendicularly from the midrib. Similarly to intact leaves, we find only a limited enrichment in the longitudinal direction in the drought leaf, and no enrichment in the sprayed leaf. The similarities in enrichment patterns between intact leaves and the experiment presented here lead us to conclude that sectioning the leaf did not significantly impact the leaf hydraulics itself over the duration of our experiment.

**Table 2:** Isotope composition, δ^18^O (‰) and δ^2^H (‰), for the lamina water at three positions along the radial direction (midrib to edge) and the longitudinal direction (bottom to tip) using the maps shown in Figure 1. n is the number of data points considered and the standard error is given in parenthesis. Overall, enrichment in the longitudinal direction is inconsistent, whereas a strong progressive enrichment is found in the radial direction in both leaves.

|  | Misted |  |  |  |  |  | Control |  |  |  |  |  |
| --- | --- | --- | --- | --- | --- | --- | --- | --- | --- | --- | --- | --- |
|  | Radial |  |  | Longitudinal |  |  | Radial |  |  | Longitudinal |  |  |
| | $\delta^{18}\text{O}$ | $\delta^2\text{H}$ | n | $\delta^{18}\text{O}$ | $\delta^2\text{H}$ | n | $\delta^{18}\text{O}$ | $\delta^2\text{H}$ | n | $\delta^{18}\text{O}$ | $\delta^2\text{H}$ | n |
| Midrib \| Bottom | 31.0 (3.2) | 81 (5) | 5 | 49.1 (4.8) | 132 (17) | 6 | 43.3 (3.6) | 82 (13) | 5 | 55.6 (5.0) | 123 (17) | 5 |
| Middle | 41.0 (3.9) | 102 (12) | 6 | 36.9 (3.9) | 91 (15) | 5 | 68.9 (11.7) | 162 (40) | 5 | 70.0 (9.6) | 167 (31) | 5 |
| Edge \| Tip | 52.6 (3.3) | 146 (13) | 5 | 36.9 (5.0) | 100 (8) | 5 | 94.2 (14.9) | 218 (37) | 5 | 80.7 (22.3) | 172 (62) | 5 |

Moreover, a thorough comparison of excised and intact leaves of *Lithocarpus edulis* under high evaporative demand (Miyazawa et al., 2011) concluded that the excised leaves had the same photosynthetic rate, stomatal conductance and transpiration rate as the intact leaves for up to 12 hours after being detached. The leaves of *Colocasia esculenta* are much larger than that of *Lithocarpus edulis*, allowing them to buffer the increased water loss from the excision more easily. Finally, none of our experiments lasted more than 8 hours, most of them lasting only 4 hours. We are therefore confident that the results obtained on excised leaves are a valid representation of the behavior of intact leaves placed in high water-stress conditions.

Finally, the experimental conditions might not be accurately describing field conditions. Indeed, the experiment represents wet leaves under relatively high temperature and incoming radiation. Since dew tends to form in the early hours of the day, often before sunrise, the lab conditions are therefore not representative of the majority of the time the leaf will be wet. However, dew tends to form preferably under clear skies, leading to high incoming radiation once the sun comes up. The goal of this study was to test the effect of leaf wetness on leaf transpiration, which is maximized when the sun is out. To amplify leaf wetness effects, we maintained the same temperature and radiation settings and kept the leaf wet for multiple hours. The results obtained are therefore likely an upper bound on the potential effect of leaf wetness on leaf water status.

## 5 Conclusion

In this study, we used the highly hydrophobic leaves of *Colocasia esculenta* to look at the impacts of dew water droplets deposition on the leaf water and isotope balance. We found that *Colocasia esculenta* plants do not have the capability to uptake dew water directly from the leaf surface. However, we found a significant decrease in transpiration by c. 30% and an increase in leaf water potential for dew-wetted leaves. We also highlighted the opposite effects of foliar uptake on leaf isotopes, which enriches leaf water in heavy isotopes, and transpiration suppression, which depletes it. Because both effects are of similar magnitude, taking both processes into accounts is crucial to properly interpret field data of foliar uptake.

More experiments are now required to understand the effects of transpiration suppression on different species and for a range of leaf shapes and sizes. Introducing stable isotopes of water in a model of leaf energy and water balance could help to interpret the competing effects of foliar uptake and transpiration suppression, give a new insight into non-steady-state transpiration, and improve the general understanding of the interaction of leaves with their environment.

## Acknowledgments

CGS and KKC acknowledge the financial support of NASA Headquarters under the NASA Earth and Space Science Fellowship Program - Grant 14-EARTH14F-241 - and of a Mary and Randall Hack ‘69 Graduate Award and the Science, Technology, and Environmental Policy Fellowship from the Princeton Environmental Institute. CGS thanks Missy Holbrook and the department of Organismic and Evolutionary Biology at Harvard University for hosting her during part of this work.

